# Investigating the topological motifs of inversions in pangenome graphs

**DOI:** 10.1101/2025.03.14.643331

**Authors:** Sandra Romain, Siegfried Dubois, Fabrice Legeai, Claire Lemaitre

## Abstract

**Background:** Pangenome graphs are increasingly used in genetic diversity analyses because they reduce reference bias in read mapping and enhance variant discovery and genotyping from SNPs to Structural Variants. In pangenome graphs, variants appear as bubbles, which can be detected by dedicated bubble calling tools. Although these tools report essential information on the variant bubbles, such as their position and allele walks in the graph, they do not annotate the type of the detected variants. While simple SNPs, insertions, and deletions are easily distinguishable by allele size, large balanced variants like inversions are harder to differentiate among the large number of unannotated bubbles and remain underexplored in pangenome graph benchmarks and analyses.

**Results:** In this work we focused on inversions, which have been drawing renewed attention in evolutionary genomics studies in the past years, and aimed to assess how this type of variant is handled by state of the art pangenome graph pipelines.

We identified two distinct topological motifs for inversion bubbles: one path-explicit and one alignment-rescued, and developed a tool to annotate them from bubble-caller outputs. We constructed pangenome graphs with both simulated data and real data using four state of the art pipelines, and assessed the impact of inversion size, genome divergence and variant density on inversion representation and accuracy.

**Conclusions:** Our results reveal substantial differences between pipelines in simulated graphs, with some inversions either misrepresented or lost. In addition, recovery rates are strikingly low in real human datasets, highlighting major challenges in analyzing inversions through pangenomic approaches.

## Background

The recent increase in the number of high-quality assembled genomes for a same species call for a switch in how we represent the genomic reference in genetic diversity analyses, from a single linear reference genome to variation and pangenome graphs [1, 2]. The interest of pangenome graphs resides in their representation of genomic variation between multiple genomes, allowing a diminution of reference bias when mapping reads, thus improving variant discovery and genotyping for the whole range of variant sizes and types [3, 4]. The current tools for pangenome graph construction rely on genome alignments, either by iterative pairwise alignments from the reference genome for Minigraph [5] and its derived pipeline Minigraph-Cactus [4], by reference-free pair-wise alignments for PGGB [6], or by tree-guided alignments for Progressive Cactus [7]. The standard pipeline in pangenome-graph based variant studies, performed in recent studies such as on chicken [8], wild grape [9] and human [10] genomes, comprises the construction of a graph from a collection of genomes, the identification of variants in the graph and their genotyping in other individuals or populations from read mapping onto the graph. The types of represented variants in the graphs differ between the tools, as Minigraph only aims to represent Structural Variants (SVs), typically variants that are larger than 50 bp, while the other tools also represent smaller variants in their graphs. Variant identification in pangenome graphs consists in analysing the graph topology to find topological motifs called *bubbles*. Bubbles are formed when (at least) two genome sequences diverge in the graph due to sequence differences, and meet again further on after the differing portion. Ideally, a SNP is represented in the graph by a small bubble where each allele carried in the input genomes forms a diverging path through a node of length 1 bp, while a SV is represented by a larger bubble where at least one of the diverging paths has a length of at least 50 bp. Pangenome graphs also contain more complex bubbles such as nested bubbles, which can be the result of nested variants (*e.g.* small variants inside an insertion). The main tools for bubble detection are *vg toolkit* [10], *BubbleGun* [11] and *gfatools* [12], the former being the most used one as it is included in Minigraph-Cactus and PGGB pipelines. Although this tool reports essential information on the variant bubbles detected in the graphs (*e.g.* position, alleles), it does not annotate the type of the detected variants. Without annotation of the bubbles’ variant category, it is difficult to analyze the number, type and size distribution of the SVs represented in a pangenome graph. SNP bubbles can be directly extracted as they have distinctive and exclusive allele path sizes of 1 bp. Likewise, bubbles having one of their allele paths with a size of 0 bp, a distinctive feature of insertions and deletions, can also be confidently annotated. However, all the remaining bubbles, in particular those with large sized paths, are more challenging to confidently annotate as a specific SV type, as their path size may correspond to more than one SV type. The large size of SVs makes them also more likely to contain nested bubbles, which is not that straightforward to represent in a VCF format. Vcfwave, from the vcflib suite [13], can re-align the allele sequences of the large bubbles reported in the *vg deconstruct* VCF to break down all their nested bubbles into a series of insertions, deletions and SNPs. Additionally, two recent works have deviated from this concept of bubble to find variants. The first one focused on a specific topological motif generated by inverted repeats or hairpin-like structures [14]. The second one, PanTree, introduces a novel definition of variant in a pangenome graph, allowing a more exhaustive detection of small variants embedded in larger structures and in particular variants in non-reference loci. But this tool has only been applied to Minigraph-Cactus graphs for the moment and was evaluated and compared to the bubble approach only for SNPs [15].

Among SVs, inversions happen to be less frequent in genomes than other types of variants such as insertions and deletions, but can reach particularly large sizes. In the human genome, a recent study estimated that inversions account for 0.4% of the diploid genome, with a median size of 17.6 kbp and a maximum size over 5 Mbp [16]. Nevertheless, inversions have an exceptional potential to promote adaptation and speciation. When in heterozygous state, they can locally reduce the effective rate of recombination in genomes, which can lead to the spread of adaptative allele combinations in populations or to genetic isolation through mutation accumulation [17–19]. Large inversions in particular, potentially including more genes, may have a higher probability of capturing non neutral allele combinations [20]. However, they are known to be more challenging to characterize and genotype than other types of SVs (through sequence based approaches) due to complex and often repetitive genomic context [21, 22].

While a general improvement of structural variant analysis using pangenome graphs has been reported, inversions are often overlooked in pangenome graph benchmarks and analyses. In two recent benchmarks of human [10] and bovine [23] pangenomes respectively or in the analyses of the wild-grape [9] and chicken[8] pangenomes, SVs were either evaluated as a whole or only insertions and deletions were reported. Only the Hickey et al. benchmark paper [4] reports some inversion counts, but only for the *Drosophila* pangenome. This illustrates the lack of tools able to distinguish complex variants such as inversions among graph bubbles.

Because identifying all types of variants is a crucial component to make the most out of pangenome graphs, there is a need to assess how inversions are handled by current state of the art pangenome graph pipelines (construction and analysis tools). In this paper, we explore how inversions are represented in pangenome graphs, and we define two different expected topological motifs for these variants. To explore the factors impacting their representation and thus the ability to detect them in these graphs, we generated pangenome graphs from both simulated genome sets and real human datasets using several state-of-the-art pipelines. We then annotated these motifs among variant bubbles identified by standard bubble callers and compared their distribution across pangenome pipelines. We show that each pipeline produces distinct patterns of inversion representation and that inversions are not always encoded as explicit alternative paths, sometimes requiring local realignment to be recognized. Finally, we introduce INVPG-annot, a tool that automatically annotate, among the variant bubbles identified in a pangenome graph, those that are likely to represent inversions.

## Results

### Expected topologies of inversions

In a pangenome graph, the nodes are labeled with genomic sequences, and the bidi-rected edges between nodes represent the adjacency and reading orientation (*forward* or *reverse*) of the sequences in genomes. In such a graph, a shared sequence between several genomes at a given locus is represented as a single node that is traversed by several paths, while a genomic variant is represented by distinct nodes connected in a particular topological motif, called a *bubble*. A bubble is a connected sub-graph where two nodes, referred as a source node and a sink node, are linked by (at least) two divergent paths – one of which (at least) is not empty. This structure can represent a single variant or several overlapping or nested variants. All distinct paths between the source and sink nodes represents all possible alleles or allele combinations of the underlying variants.

Contrary to other SV types (*e.g*. insertions, deletions), an inversion does not imply a modification of sequence content between alleles. As such, in the simplest event of inversion (without any inner sequence variation), we can expect both allele sequences to be represented as a single node *n_I_* in the graph, that is traversed in opposite directions (forward and reverse) by the ancestral (*p_A_*) and inverted (*p_I_*) allele paths (Fig. 1a). As inversions can cover large portions of the genome, it is most likely that inversion loci contain inner smaller variants (*e.g*. SNPs, indels). If the graph building algorithm represents small variants in the graph, those will break the inversion locus into multiple nodes and the list of nodes traversed by *p_A_* and *p_I_* will differ (a simple example with one SNP is illustrated in Fig. 1b). Nevertheless, we expect the cumulative length of the nested variants to be much smaller than the total length of the inversion, allowing for the existence of nodes that are common and traversed in opposite directions between *p_A_* and *p_I_*. In both simplistic and realistic cases, the inversion event should be explicitly represented by the paths of the bubble, and so we define this topology type as a “path-explicit” inversion representation.

**Fig. 1.**
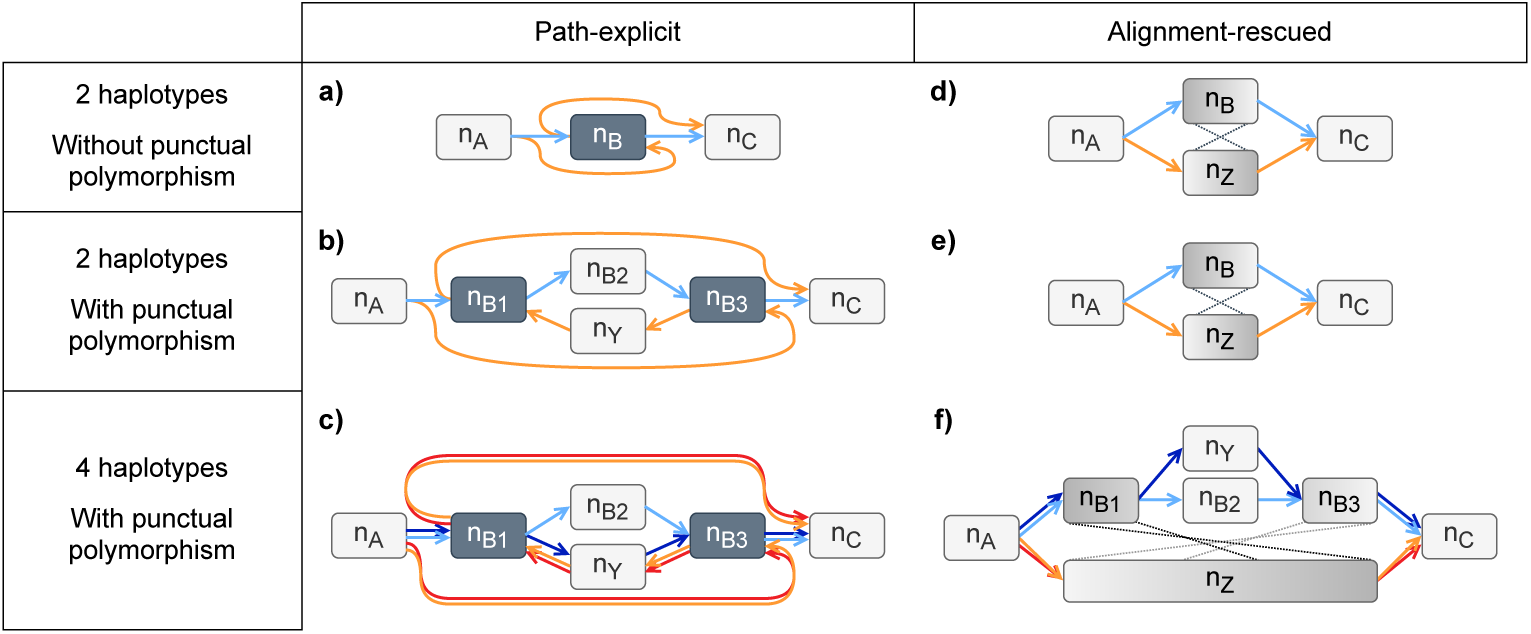
Illustration of the expected types of bubble topologies for inversions in pangenome graphs. Arcs of the graphs are colored according to the presence (orange and red) or absence (blue) of the inversion. The examples on the left illustrate cases where an inversion is explicitly represented in the paths of a bubble (”path-explicit”), with the dark grey colored nodes being the ones that are traversed in both directions **(a, b, c)**. The examples on the right illustrate cases where the two alleles of an inversion are represented by distinct nodes and paths (”alignment-rescued”), with grey-shaded nodes indicating pairs of nodes that are the reverse complement of each other **(d, e, f)**. For each expected topology type, different levels of complexity of the graph are presented: a first level with two haplotypes and no SNP polymorphism inside the inversion **(a, d)**, a second level with two haplotypes and a SNP between the two inversion alleles (**b, e**), and a third level with four haplotypes, one of which (*dark blue*) carries the reference allele of the inversion and the alternative allele of the SNP (**c, f**).

However, if an inversion is not identified, meaning the homology between the dif-ferent alleles of the inversion is not detected, either during the genome alignment step or during the PG graph construction, both ancestral and inverted alleles could be represented as unrelated sequences in the graph. In the simplest cases with two hap-lotypes, the locus could be represented by two entirely disjoint paths, one traversing a node *n_B_* carrying the ancestral version of the sequence, the other traversing a node *n_Z_* carrying the inverted version of the sequence (Fig. 1 d,e). In this case, in order to identify the inversion, one path sequence must be compared in its forward orientation to the reverse complement of the other path sequence, and so we define this topology type as an “alignment-rescued” inversion representation.

We developped INVPG-annot, an automated method to identify, among the bubbles found in a pangenome graph, the ones that correspond to these inversion topologies. The method is composed of three main steps, as shown in Fig. 2. The first step consists in selecting potential inversion bubbles based on the size of their alleles, in order to limit the amount of bubbles to analyze. Then in the second step, for each selected bubble, inversion signals are searched by comparing the paths of the different alleles and identifying nodes traversed in opposite directions in different alleles. We compute a path-based inversion signal score *cov_path_*reflecting the fraction of the reference allele’s length covered by path-based inversion signals. If *cov_path_*is greater than a user-defined threshold (*min_cov_*, with default value of 0.5), the bubble is annotated as a path-explicit inversion. If not, we then apply the third step, and look for inversion signals in allele sequence alignments. We align all alternative alleles sequences on the reference allele sequence using minimap2 [24] and compute an alignment-based inversion signal score *cov_aln_*reflecting the fraction of the reference allele’s length covered by reverse alignments. If *cov_aln_* is greater than the *min_cov_* threshold, the bubble is annotated as an alignment-rescued inversion (see the details in Methods).

**Fig. 2.**
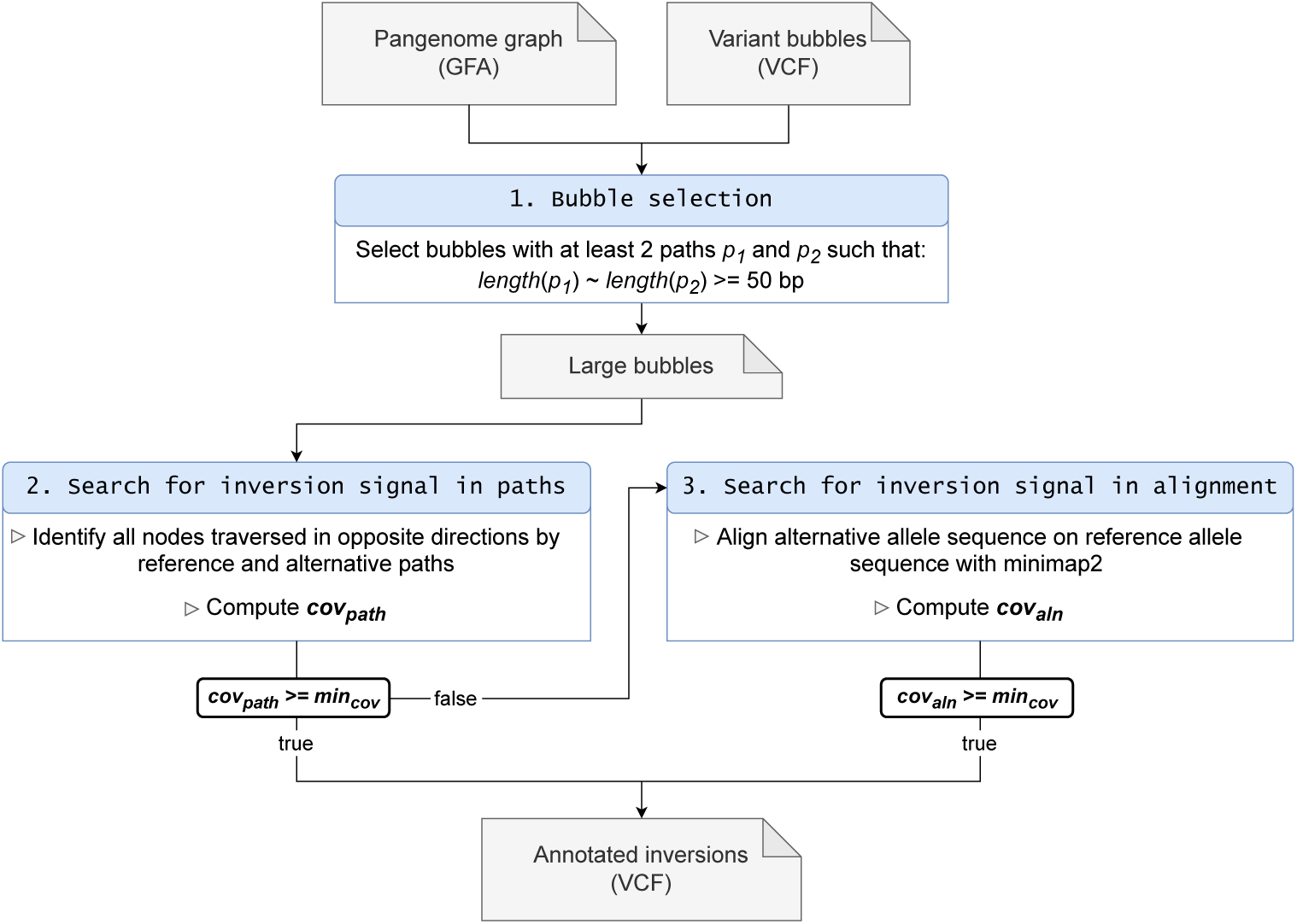
Illustration of the inversion annotation method INVPG-annot. The method takes as input a variant bubble file in VCF format and a pangenome graph in GFA format. The first step of the method consists in selecting potential inversion bubbles based on their allele path sizes. Selected bubbles are then searched for path-based and alignment-based inversion signals. Bubbles annotated with an inversion signal score greater than a used-defined threshold are output in VCF format.

### Investigating inversion bubble topologies in simulated graphs

We generated several simulated haplotype sets from the CHM13 human chromosome 21 with well-controlled inversion variants in order to investigate how inversions are represented in pangenome graphs of various complexities. The inversions we included in these simulated haplotypes were non-overlapping, with randomly positioned coordinates, and with sizes ranging from 50 bp to 1 Mb, equally distributed in four size classes, 50-1,000 bp, 1-5 Kb, 5-100 kb and 100-1,000 Kb (see details on size distribution in Methods). We generated several haplotype sets, by varying the inversion density in the simulated haplotypes (total inversion number ranging from 40 to 140), the number of haplotypes in the graph (2 or 10) and the level of SNP divergence between haplotypes (SNP percentage varying from 0 to 5%).

For each simulated haplotype set, we built their pangenome graphs and called the bubbles with the main pangenome graph pipelines, namely Progressive Cactus[7] (referred to as Cactus in the following Results), Minigraph[5], Minigraph-Cactus[4] (MC) and PGGB[6] (see Methods). We then applied our method to look for inversion signals in these bubbles and measured, for each dataset-pipeline combination, a recall value, corresponding to the percentage of the simulated inversions that are effectively annotated as inversion bubbles (see Methods for details on the comparison of inversion calls).

### Most inversions are represented as bubbles in the graphs

We show in Table 1 the graph and bubble statistics obtained for the four pangenome pipelines with the genome set composed of 2 haplotypes with 100 inversions and a SNP divergence of 1%. All four pipelines produced graphs of total size varying from 28.28 to 29.78 Mb, which is slightly larger than the size of one haplotype (28 Mb). In terms of bubbles, except for Minigraph which is intended to represent only large variants, all other three pipelines produced hundreds of thousands of bubbles. For the Cactus and PGGB graphs, the number of bubbles that pass our method’s filter on bubble size is close to the 100 expected (92 large bubbles). For MC and Minigraph graphs, the number of large bubbles detected is fewer than expected, with respectively 86 and 75 large bubbles. For all four graphs, all or almost all of the large bubbles are annotated as inversions. Overall, between 75 and 92% of the simulated inversions were represented as inversion bubbles in all four pangenome graph pipelines. More precisely, we obtained the best recall value of 92% with the Cactus pipeline, followed by MC with 82% and PGGB with 89%. The smallest recall value was observed with Minigraph pipeline with a value of 75%.

**Table 1.** Graph and bubble statistics of the pangenome graphs obtained for the 2-and 10-haplotype simulated genome sets with 100 inversions and 1% of SNP polymorphism, for each pangenome graph pipeline used. All haplotypes in these genome sets have a size of 28 Mb. The number of bubbles indicates the total number of bubbles reported by the pipeline bubble caller. Large bubbles are those selected at the first step of INVPG-annot. Inversion bubbles are those annotated as inversions by INVPG-annot. True positives (TPs) are inversion bubbles that pass the recall overlap constraints. The Recall column indicates the percentage of simulated inversions (among the 100) that have a significant reciprocal overlap with at least one inversion annotated bubble. The number of TPs and recall value can differ in case of redundant TPs. (”MC”: Minigraph-Cactus).

| # Hap. | PG pipeline | Graph size (Mb) | # Bubbles | # Large bubbles | # Inversion bubbles | # TPs | Recall (%) |
| --- | --- | --- | --- | --- | --- | --- | --- |
| 2 | Cactus | 28.28 | 277,961 | 92 | 92 | 92 | 92 |
|  | Minigraph | 28.32 | 75 | 75 | 75 | 75 | 75 |
|  | MC | 29.78 | 322,643 | 86 | 84 | 82 | 82 |
|  | PGGB | 28.43 | 275,670 | 92 | 90 | 89 | 89 |
| 10 | Cactus | 28.57 | 549,460 | 167 | 162 | 161 | 90 |
|  | Minigraph | 28.32 | 77 | 77 | 76 | 76 | 76 |
|  | MC | 31.39 | 550,516 | 92 | 88 | 86 | 86 |
|  | PGGB | 28.73 | 548,125 | 115 | 94 | 92 | 92 |

Notably, we did not observed any false positive among the inversion annotated bubbles. Only one and two inversion bubbles for PGGB and MC, respectively, were not counted as True Positive (TP), but we do not classify them as false positives because they overlap with simulated inversions, only the overlap is below the required threshold of 50% reciprocal overlap (these bubbles are referred to as imprecise in Supplementary Table 1).

We observed no major difference in recall at other divergence level (0.1 or 5%, Supplementary Table 1). We did observe a curiously low number of annotated inversions (7 and 28) and even lower recall (4% and 24%) with Minigraph and MC graph at 0% SNPs, but interpreted it as an artefact or bug of the software from the Minigraph pipeline, which occurs only for this unlikely rate of divergence, and which we do not expect to occur under realistic levels of genomic divergence. Increasing the number of haplotypes to 10 in the graphs at 1% SNP polymorphism also did not seem to impact greatly the inversion recall, with at most a 4 point percentage difference with the MC graph (Tab. 1).

### Two inversion topologies can be found in the graphs

Bubbles annotated as inversions were divided in two types: (i) “path-explicit”: the signals of inversion can be directly identified in the graph (ii) “alignment rescued”: the inversion has been detected after alignment of the bubble path sequences. As a result, we observed that the pangenome pipelines differ in the bubble topology they use to represent inversions (Fig. 3A). In the graphs built with the 2-haplotype and 100-inversion genome set at 1% divergence, all inversion bubbles detected in the Cactus graph are path-explicit. In Minigraph and MC graphs, most of the annotated bubbles are also path-explicit, with only 3 and 10 inversion bubbles with an alignment-rescued topology, making up for 4 and 12% of the annotated bubbles in those graphs, respectively. On the other hand, the two inversion bubble topologies are almost perfectly equally represented in the PGGB graph. Interestingly, we found that the inversion topology in PGGB is strongly linked to the inversion size, as all path-explicit inversion bubbles represent inversions larger than 17 kb, and all alignment-rescued ones represent inversions smaller than this threshold (Fig. 4B). Even when the inverted segment contains no single nucleotide polymorphism, all inversions smaller than 17 Kb are still represented as two independent nodes in the PGGB graph, one being strictly the reverse complement of the other (alignment-rescued topology). Notably, we observed an increase in the proportion of inversions bubbles with the alignment-rescued topology when increasing the SNP divergence to 5% in Minigraph and MC graphs, with respectively 50 and 51 alignment-rescued inversion bubbles, accounting for more than half of the annotated bubbles in these graphs (Fig. 4A).

**Fig. 3.**
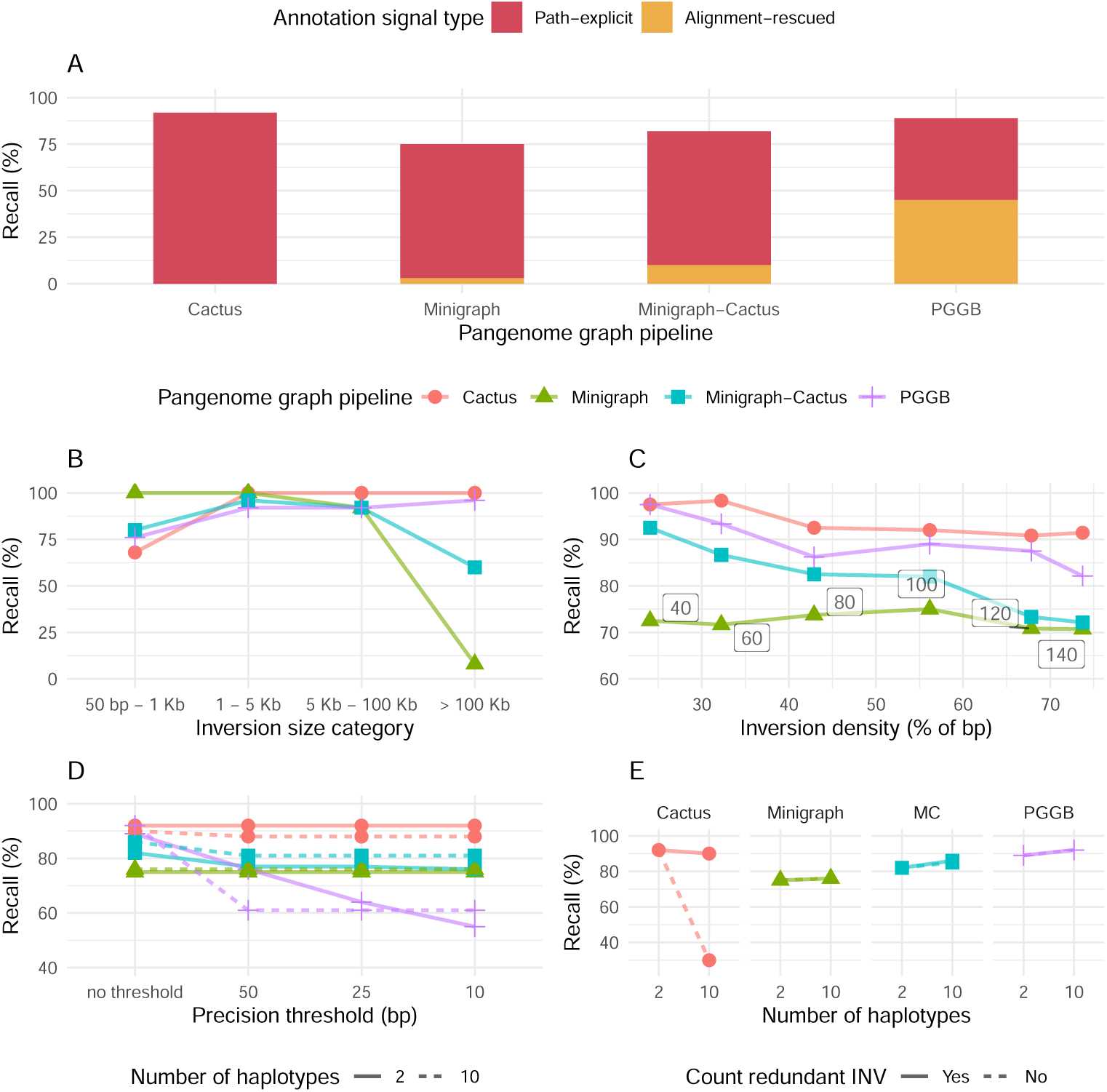
Evaluation of inversion annotation on the pangenome graphs built with the four pipelines from the simulated datasets. The main simulation scenario contains 100 inversions in 2 haplotypes at 1% SNP polymorphism. (A) Recall of inversion annotation as calculated by the percentage of simulated inversions overlapped by at least one bubble annotated as inversion. Each recall bar is divided into two colors: red for inversions obtained from path-explicit annotations and yellow for alignment-rescued annotations. (B) Variation of the recall depending on inversion size category. (C) Variation of the recall depending on the inversion density in the haplotypes (calculated as the percentage of inverted base pairs). The numbers in the boxes indicate the number of inversions simulated at each point of density level. (D) Variation of the recall when increasing the level of bubble boundary precision to consider an inversion bubble as a True Positive, for datasets with 2 (straight line) and 10 (dashed line) haplotypes. To be counted as a True Positive, the bubbles must have a reciprocal overlap of at least 50% with the true inversion coordinates and both breakpoints are less than X bp offset from the true coordinates, X taking the values 10 bp (most precise), 25, 50 bp and infinity (least precise without any absolute threshold applied). (E) Recall counting redundant inversion bubbles (straight line) and recall dismissing redundant inversion bubbles (dashed line) depending on the number of haplotypes in the graphs. The line colors and datapoint shapes in B-E correspond to the pangenome graph pipeline used.

**Fig. 4.**
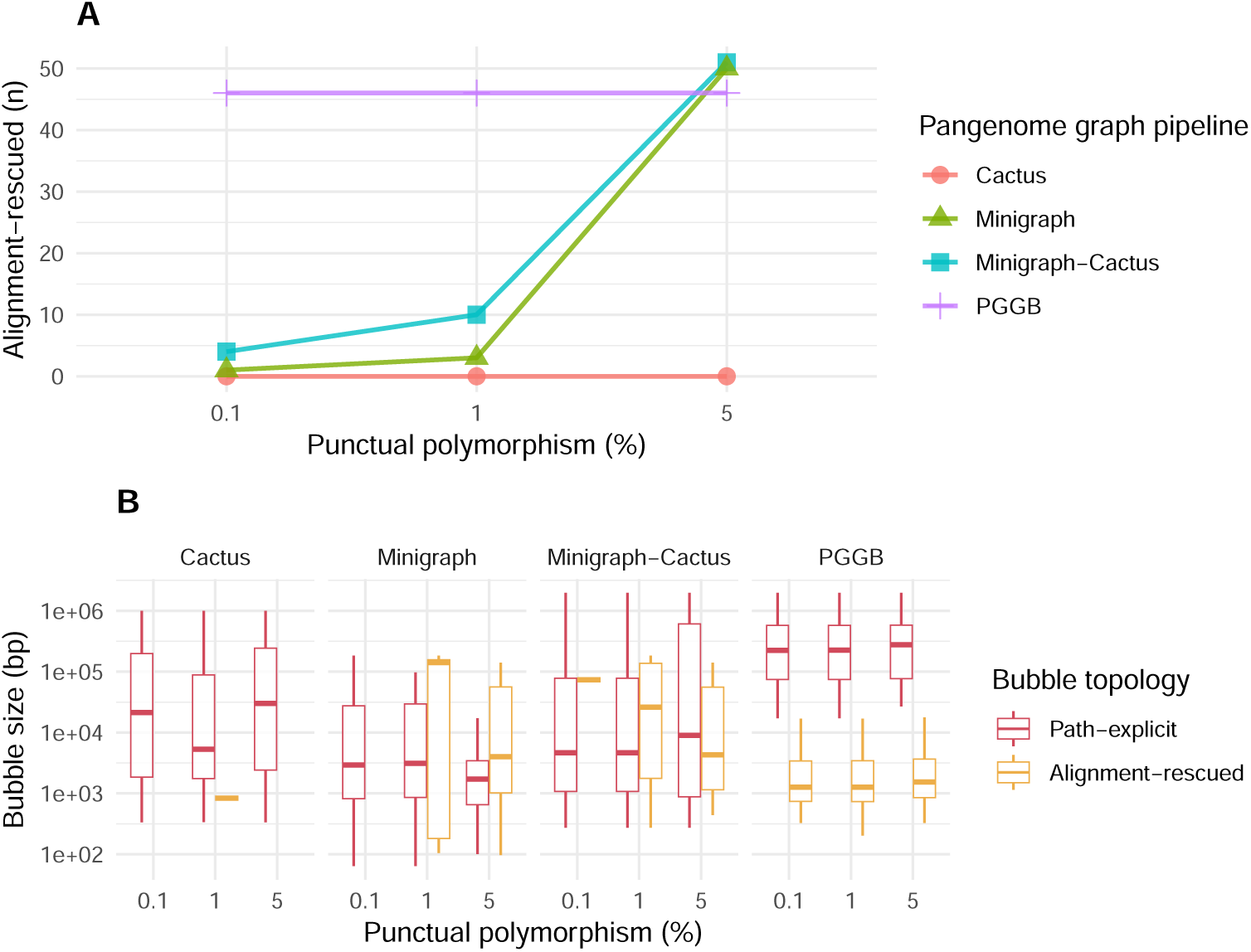
Topology of the true positive inversions annotated in the pangenome graphs built with the 4 pipelines from the simulated datasets with 100 inversions in 2 haplotypes depending on the level of SNP polymorphism. (A) Number of true positive alignment-rescued inversion bubble in the four pangenome graph pipeline. (B) Size distribution of the true positive path-explicit (red) and alignment-rescued (yellow) inversion bubbles.

### Inversion size affects inversion recall

To determine if the size of the inversions could impact their representation in pangenome graphs, we calculated the recall for each inversion size category in the graphs built with the 2-haplotype and 100-inversion genome set (Fig. 3B). All four pangenome graph pipelines produced graphs yielding top recall values (92 to 100%) for intermediate size categories of inversions (between 1 and 100 Kb). However, the four pipelines showed a drop of inversion recall for either the smallest (Cactus and PGGB) or the largest inversions (Minigraph), or both (Minigraph-Cactus). The most dramatic drop of inversion recall was for inversions larger than 100 Kb in the Mini-graph graph, with only 8% of these large inversions being recovered. In Cactus and PGGB graphs, the recall of inversions below 1 Kb dropped to 68 and 76%, respectively. Interestingly, the recall trend in MC graph showed a combination of Minigraph and Cactus trends, with drops for both the smallest and largest size ranges, though more moderated than those of Minigraph and Cactus alone.

### Inversion density in the pangenome impacts the recall

We also evaluated whether the total number of inversions simulated in the haplotypes can impair the representation of each individual inversion, keeping all inversions non-overlapping and spaced at least 1 Kb apart. In Fig. 3C, we show the obtained inversion recalls for varying inversion densities. We calculated inversion density as the percentage of base pairs in a haplotype covered by an inversion (*i.e.* a density of 50% means that half of the haplotype length is covered by inversions). With the 2-haplotype graphs at 1% SNP divergence, we observed a general trend of decreasing recall when increasing the inversion density for all but Minigraph’s graph. The graphs most impacted by the inversion density were those of MC and PGGB, going from a 92.5% and 97.5% recall at 24% density (40 inversions per haplotype) to a 72.1% and 82.1% recall at 74% density (140 inversions per haplotype), respectively. In the case of Cactus graphs, inversion recall dropped by less than 10 percent point, from 97.5% at 24% density to 91.4% at 74% density.

### Bubble precision and redundancy

In addition to the inversion recall, we evaluated the quality of inversion representation in pangenome graphs by the precision and redundancy of their bubbles. In Figure 3D, we show the variation in recall values when filtering annotated inversions based on their breakpoints’ precision, using an increasingly stringent threshold, for both 2- and 10-haplotype graphs at 1% SNP divergence. On our simulated data, all PG pipelines but PGGB show high precision for inversion bubbles. Cactus and Minigraph pipelines construct graphs with the most precise inversion bubbles, of which either breakpoint positions are at most 10 bp offset from the true positions. Most of inversion bubbles in MC graphs also have a precision of 10 bp or under, as for both 2- and 10-haplotype graphs, the recall drops by only 6 point percentage when applying the 10 bp precision thresholds. On the other hand, PGGB graphs show the least precise inversion bubbles, with recall values of only 55% and 61%, respectively, when applying the 10 bp precision threshold with 2 and 10 haplotypes. This means that half of the inversions represented in PGGB graphs and annotated by our method have bubble boundaries located more than 10 pb away from the true coordinates of the inversions.

Since bubbles can be nested within each other, we may annotate inversion signals in several overlapping bubbles. A single simulated inversion can then be represented in several bubbles, which we refer to as redundant. We evaluated the inversion bubbles redundancy by re-calculating the inversion recall dismissing all true positive (TP) redundant bubbles. While Minigraph, MC and PGGB graphs were almost bare of redundant bubbles for both the 2- and 10-haplotype sets and thus showed no decrease of recall when dismissing TP redundant bubbles, the Cactus graph showed a multiplication of redundant bubbles for the 10-haplotype set, with 132 redundant inversion bubbles making up for more than 80% of the annotated inversion bubbles, and a decrease of 60 points % of the recall when dismissing TP redundant bubbles (Fig. 3E, Sup.Tab. 2).

**Table 2.** Bubble statistics obtained for the combined chromosomes 7 and X real human haplotype sets, for each pangenome graph pipeline (”MC”: Minigraph-Cactus). True positives (TP) are validated inversion bubbles (*i.e.* annotated bubbles passing the recall constraints). The Recall column indicates the percentage of inversions from the truth set (among the 41) that have a significant reciprocal overlap with at least one inversion annotated bubble.

| PG pipeline | # Bubbles | # Large bubbles | # Inversion bubbles | # TP | Recall (%) |
| --- | --- | --- | --- | --- | --- |
| Cactus | 1,512,364 | 4,111 | 14 | 4 | 9.8 |
| Minigraph | 4,620 | 95 | 29 | 22 | 53.7 |
| MC | 1,042,944 | 1,707 | 25 | 22 | 53.7 |
| PGGB | 1,024,316 | 2,675 | 50 | 8 | 19.5 |

### Inversion bubble annotation in real pangenome graphs

Finally, we investigated how real inversion variants are represented in pangenome graphs constructed with real human haplotype sequences. Based on the inversion polymorphism study of [16], we selected four human individuals, genotyped in this study and with an available diploid genome assembly from the HPRC draft human pangenome [10]. We selected two inversion-rich chromosomes, chromosomes 7 and X, harbouring in total 41 inversions, with a 0/1 or 1/1 genotype for at least one of our four selected individuals (see Methods). In this inversion set, which we use as a truth set, inversions range from 1.1 kb to 1.7 Mb, with a median size of 32 kb. For each chromosome, we built graphs with nine human haplotypes (the four selected diploid individuals and the GRCh38 reference sequence) and called variants with the four pangenome pipelines. We then applied our method, INVPG annot, to annotate inversion bubbles and computed a recall value corresponding to the percentage of the 41 inversions from the truth set that are effectively annotated as inversion bubbles.

Compared to the results with simulated haplotypes, we obtained a much lower recall of inversion annotation for all pangenome pipelines on the combined two chromosomes, with no more than 9.8% to 53.7% of the expected inversions that were found among the called bubbles (Table 2). Cactus showed the lowest recall of 9.8%, followed by PGGB with 19.5%, and Minigraph and MC with the highest recall of 53.7%. Contrasting the TP inversion bubbles by size category, Minigraph and Minigraph-Cactus showed the highest recall values between 63 and 75% for the 1 to 5 Kb and 5 to 100 Kb inversions (Figure 5A). All pipelines produced graphs with poor recall for inversions larger than 100 Kb, from 0% recall with Cactus and Minigraph to 20% recall for PGGB. Interestingly, the sets of inversions annotated on the graphs showed differences between the pipelines, with some inversions being annotated in only one graph (Figure 5B). Overall, 28 inversions out of the 41 true inversions (68%) were detected in at least one pipeline’s graph, but only 2 were commonly detected in all four graphs and 7 were detected in only one graph. Out of all TP inversion bubbles for the different graphs, only 1 had an alignment-rescued topology in the Minigraph and Cactus graphs (amounting to 4.5 and 12.5% of the TP inversion bubbles, respectively), and 5 in the PGGB graph (amounting to 41.6% of the TP inversion bubbles) (Figure 5C).

**Fig. 5.**
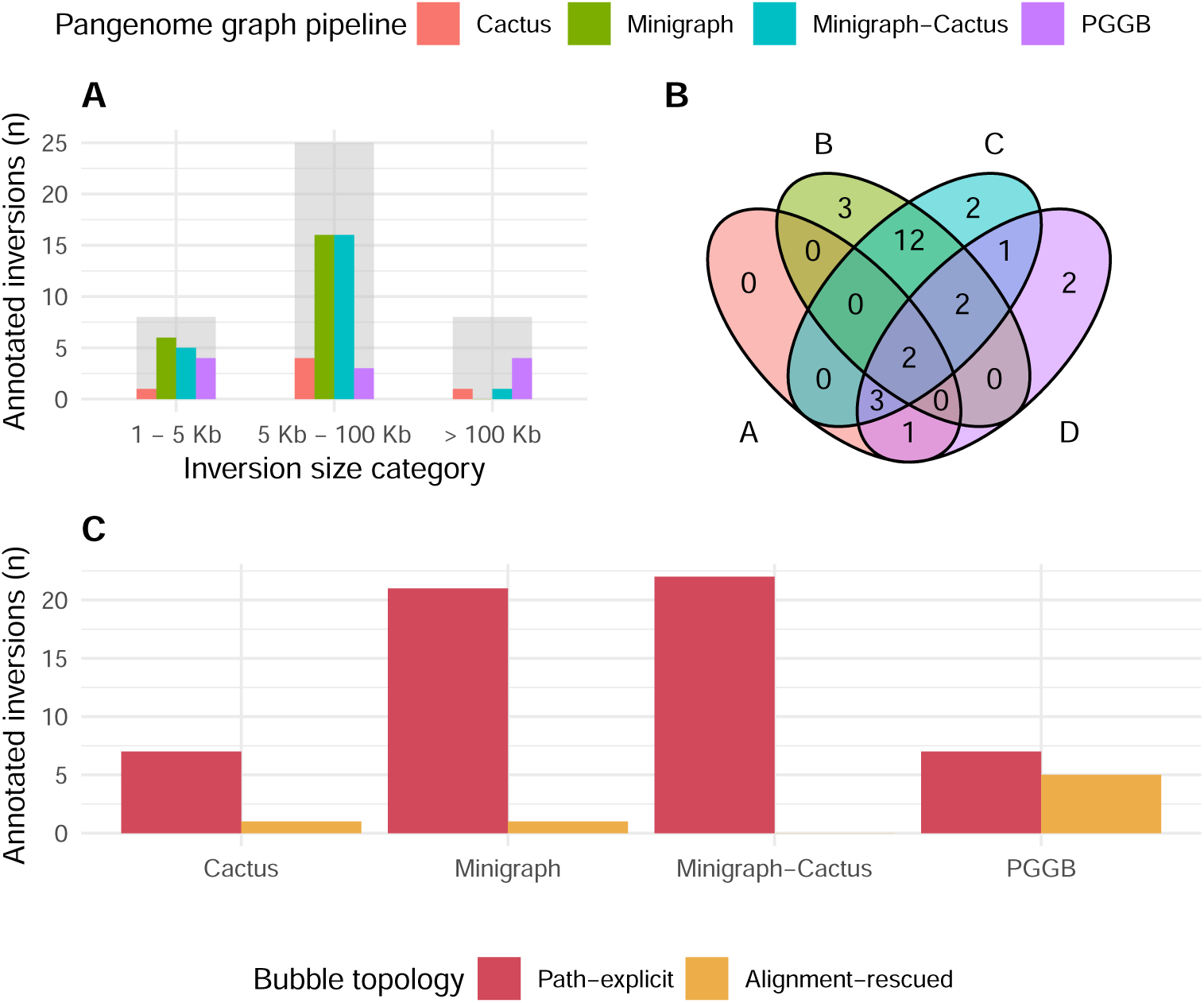
Inversions annotated in the pangenome graphs built with the 4 pipelines from the real human dataset (9 human haplotypes of chromosomes 7 and X). Only the true positive inversions by comparison to the real inversion truth set are shown. (A) Number of true positive inversion bubbles annotated in the pangenome graphs depending on inversion size category. The grey boxes behind show the total number of inversions of the same size categories in the truth set. (B) Venn diagram showing the intersection of true positive inversion bubbles sets between each of the 4 pangenome pipelines. The intersection between the inversion sets were evaluated from the overlap of the inversion positions on the reference genome, using the same overlap constraints as that of the recall (as described in Methods). (C) Number of true positive path-explicit (red) and alignment-rescued (yellow) inversion bubbles for each pangenome graph pipeline.

## Discussion

In this study, we investigated how inversion variants, a major class of balanced struc-tural variants, are represented in pangenome graphs. We first formalized the expected representation of inversions in ideal pangenome graphs. We identified two expected topologies, both forming bubbles but constrained by the relationships between their alternative paths: a path-explicit topology, in which the alleles traverse the same nodes in reverse-complement orientation, and an alignment-rescued topology, in which the alleles pass through distinct nodes that are reverse-complement similar. Using controlled simulated datasets, we then confirmed that most inversions are indeed represented as bubbles in graphs produced by current state-of-the-art pipelines, and that both topologies occur in practice. Building on these structural signatures, we devel-oped INVPG annot, a method that annotates bubbles detected by standard bubble callers and distinguishes inversion bubbles from the vast majority of other variant bubbles. It also reports which inversion topology is present, allowing for their precise characterization in pangenome graphs. INVPG annot is to our knowledge the first available tool to annotate inversion bubbles in pangenome graphs with a VCF output, allowing a seamless integration in classical pangenome variant analysis pipelines. It provides a fast annotation, annotating inversions in 300+ Mbp pangenome graphs in under 20 minutes when the alignment requirements of the graph bubbles are low (Supplementary Table 3).

We next assessed the accuracy of inversion representation using simulated datasets designed to capture different genomic scenarios, including variation in inversion size, number of haplotypes in the graph, SNP sequence divergence and inversion density.

Overall, most inversions were represented as bubbles, but recall ranged between 8% and 100% between the different scenarios and between the pipelines. We observed notable differences between graph construction pipelines in both the range of inversion sizes that could be accurately represented and the relative frequency of the two topologies. In particular, very large inversions (above 100 Kb) were rarely represented in Minigraph graphs, although this tool has been specifically designed to represent only large variants.

The pangenome graph pipelines also showed different patterns of inversion topology distribution, and the need to re-align bubble alleles proved to be especially relevant for inversions with a high SNP divergence for MC and of smaller size (below 17 kb) in the PGGB graphs. The impact of sequence divergence was expected: as divergence increases, homology is more frequently missed during the alignment step, leading the graph construction process to represent the alleles as unrelated nodes rather than alternative traversals of the same nodes. Consistently, in MC graphs only 10% of inversions showed an alignment-rescued topology at 1% SNP divergence, whereas this proportion increased to 50% at 5% SNP divergence. Although 5% divergence exceeds typical levels observed in human genomes, such values can be realistic in other taxa, including plants and insects, and may occur particularly within inversions themselves. Because large inversions can suppress recombination between the inverted and non-inverted forms, each orientation can accumulate lineage-specific mutations, resulting in substantially higher divergence than in surrounding genomic regions [18, 20].

In contrast, the dependence on inversion size observed in PGGB graphs is less intuitive and appears primarily driven by alignment parameterization of PGGB pipeline. We found that the threshold separating path-explicit and alignment-rescued topologies largely depends on the wfmash parameters controlling seed length and alignment chaining (parameters-s,-l, and-c). When wfmash produces long alignments spanning small inversions, these events are not recovered by seqwish, which infers variation solely from CIGAR strings. Consequently, inversions embedded within long alignments were not explicitly represented as alternative paths. After exploring multiple parameter combinations, we observed that the default values of-s,-l, and-c provided the highest inversion recall in our simulated datasets.

While inversions represented as unrelated sequences in the graph (*i.e.* with distinct nodes for the ancestral and inverted versions of the sequence) can still be recovered if there is at least one bubble containing the entire locus, they need additional alignment to be annotated, which can become computationally costly in large and complex graphs (for example, ∼35 minutes for annotating bubbles from two chromosomes in the PGGB human pangenome graph). Beyond computational cost, this representation introduces redundancy because sequences sharing a common origin and genomic position are duplicated in the graph. Such duplication may complicate downstream analyses. For instance, small variants located within these inversions raise ambiguity in both representation and genotyping: variants may appear on one path, on both paths, or be missed entirely, and read mapping may become ambiguous. Reads originating from the inverted locus may align to either path, increasing multi-mapping and reducing genotyping reliability. Overall, inversions that are not explicitly modeled as alternative traversals of the same locus should ideally be represented in a path-explicit topology to minimize redundancy and ambiguity, thereby improving the interpretability and usability of pangenome graphs.

Our analysis also revealed the prevalence of redundancy among inversion bubbles in the Cactus graphs containing a higher number of haplotypes or a higher percentage of SNPs, as well as breakpoint position imprecision in the PGGB graphs. These two properties limit the usability and reliability of annotated inversions in these graphs. One way to circumvent these limitations could be to develop specific methods to refine the breakpoints of inversions in bubbles, such as by analyzing the distribution of the inversion signal along the paths of the bubbles A notable result of our analysis is the marked discrepancy between simulated and real datasets. In controlled simulations, inversion recall consistently reached 80–90% across pipelines, whereas in human pangenome graphs built from real data it dropped to 10–54%. One possible explanation is the larger diversity of variants in real haplotypes. Unlike the relatively isolated inversions introduced in our simulations, human genomes can contain dense combinations of SNPs, indels, and other structural variants in the same regions. Such local complexity can disrupt alignment anchoring and chaining during graph construction. As a matter of fact, we observed that an overall increase in inversion density in our simulated chromosomes affected recall even when additional inversions were introduced at distant loci, suggesting that graph construction heuristics are globally influenced by variant load rather than purely local sequence context.

A second, non-exclusive explanation is that real inversions are structurally more complex than the simplified models typically simulated. Human inversions can exhibit breakpoint-associated variation, including small insertions and deletions at the break-points. These features may blur the boundaries of the inverted segment and may fragment its representation in the graph, preventing their detection as bubbles by bubble-caller tools. Additionally, human inversions are frequently flanked by large inverted repeats or segmental duplications [16], which can lead to ambiguous alignments and collapse or duplication of nodes during graph induction, further obscuring the inversion topology. The tendency of PGGB to collapse duplications into common nodes and generate more cyclic graph structures than MC likely reduces the formation of detectable bubbles; accordingly, PGGB exhibited a larger drop in recall from simulated to real data (92% to 20%) than MC (86% to 54%).

Together, these observations suggest that current graph representations capture idealized inversions more readily than biologically realistic ones. An interesting avenue for future work would therefore be to extend our motif definitions to incorporate additional structural signatures of inversions.

Another possible explanation is that some inversion-associated topologies are present in the graph but remain unreported by current bubble detection tools. These methods may rely on heuristics or require structural constraints for the bubbles, which are not always clearly described in the literature. For example, vg deconstruct does not report non-nested overlapping variants. While this limitation has little impact in our simulated datasets, where variants were generated without overlap, it may become relevant in real genomes where structural variants may intersect (for instance, if a deletion in one haplotype overlaps an inversion breakpoint). Unfortunately, we could not compare multiple bubble callers on the same graphs, because vg-deconstruct and gfatools require different graph formats (GFA with P or W lines for the former and rGFA for the latter), and BubbleGun could not be used as it does not report path traversals within bubbles. A potential extension of our approach would therefore be to search the graph directly for inversion-specific topological patterns in order to bypass the constraints of the bubble detection tools. However, directly exploring the graph in this manner may be computationally intensive and was beyond the scope of the present study.

## Conclusion

In this work, we characterized how inversions are represented in pangenome graph bubbles, identifying two main bubble topologies and confirming their occurrence in simulated and real datasets. Using these insights, we developed INVPG annot to automatically annotate inversion bubbles and distinguish their structural topology. In simulated graphs, we observed substantial differences between pipelines, with some inversions either misrepresented or lost. In addition, recovery rates are strikingly low in real human datasets, highlighting major challenges in analyzing inversions in pangenomic approaches. To our knowledge, this is the first study of how inversions are represented in pangenome graphs, providing a strong basis for the broader detection and characterization of such variants in pangenomes.

## Methods

### Simulation of genome datasets from the human chromosome 21

We based our simulations on the genomic sequence of the chromosome 21 of the CHM13 human reference genome, in which we removed the peri-centromeric and sub-telomeric regions [25]. We extracted the sequence between coordinates 17 Mb and 45 Mb, resulting in a genomic sequence of 28 Mb. This sequence constitutes the reference haplotype in all simulated haplotype sets, and was mutated to generate other haplotypes.

### Inversion sets

We defined a set of 100 non-overlapping inversions, whose coordinates were randomly positioned on this chromosome. We divided equally the set in 4 size categories, 50 bp to 1 Kb, 1 Kb to 5 Kb, 5 Kb to 100 Kb, and 100 Kb to 1 Mb. In each size category, the sizes of the 25 inversions were sampled uniformly in the corresponding size range. In addition, we ensured that all inversions were more than 1 Kb away from each other. For the analysis of the inversion density impact, we generated five more inversions sets containing 40, 60, 80, 120 and 140 inversions. We controlled the inversion positions and sizes in the same fashion as done for the 100 inversion set, sampling *n/*4 sizes per size range, with *n* being the total number of inversions in the set.

### Haplotype sets

We simulated one synthetic chromosome for each of the six sets of inversions, with additional single nucleotide mutations uniformly introduced at a 1% rate. As a result, we obtained six 2-haplotype sets each corresponding to a different inversion density level, the first haplotype being the reference chromosome sequence and the second haplotype being one of the synthetic chromosomes.

For the analysis of the number of haplotypes, we generated a set of 10 haplotypes. We first generated a set of uniformly positioned single nucleotide mutations corresponding to a 2% rate. We then generated nine different haplotypes by mutating the reference sequence with different subsamples of 50% of the single nucleotide mutation set and 50% of the 100 inversion set. We ensured that every inversion of the 100-inversion set was present in at least one of the nine haplotypes.

For the analysis of nucleotidic divergence impact, we simulated four synthetic chromosomes containing the set of 100 inversions, with additional single nucleotide mutations uniformly introduced at different rates: 0, 0.1, 1 and 5% rate. We obtained four 2-haplotype sets each corresponding to a different nucleotide divergence level, the first haplotype being the reference chromosome sequence and the second haplotype being one of the synthetic chromosomes.

### Real haplotype and inversion sets

For application on real human haplotypes, we based our haplotype set construction on the extensive inversion dataset published by [16], reporting 292 balanced inversions across the GRCh38 reference and genotyped in 44 individuals. We used the GRCh38 human reference genome, as well as the diploid genome assemblies of the four individuals NA19240, HG00733, HG03486 and HG02818 published by the HPRC [10], which were the only assemblies available among the 44 genotyped genomes. In order to limit the processing cost, we selected two chromosomes to process. We filtered the inversion dataset to remove unbalanced inversions (*i.e.* duplicated inversions) and low confidence inversion calls (without the’PASS’ tag). We finally selected the chromosomes 7 and X, of 159 Mb and 156 Mb respectively, which with 16 and 25 inversions respectively present in at least one of the four selected individuals were the most inversion-rich chromosomes in this dataset.

We ran RagTag v2.1.0 [26] to resolve the assemblies’ contigs correspondance to GRCh38 chromosomes 7 and X, and extracted the corresponding (unscaffolded) contigs in separate multi-fasta files for each of the 8 haplotypes.

### Construction of the pangenome graphs and bubble calling

For each set of haplotypes, simulated or real, we constructed four versions of pangenome graphs using Minigraph [5], Minigraph-Cactus [4], PGGB [6] and Progressive Cactus [7].

We ran Minigraph-Cactus v2.9.9 with the parameters ‘–clip 0 –filter 0‘ to retain all variants and haplotype paths in the graphs. PGGB v0.7.4, Minigraph v0.21 and Progressive Cactus v6.0.0 were run with default parameters. As it is recommended to use binarized input trees with the more recent versions of Progressive Cactus, we wrote arbitrary trees with uniform branch length to ensure they have the right structure, keeping haplotypes of single individuals closest together in the trees. We recon that the trees should have minimal impact on human and simulated data experiments as genomes are close. We then converted the resulting HAL file of Progressive Cactus to a VG graph format using the *hal2vg* tool with the‘–noAncestors‘parameter. *Vg convert* [3] v1.65, was then used with the ‘-f-W‘ parameters to obtain the Minigraph-Cactus and Progressive Cactus graphs in GFA 1.0 format.

Variant bubbles were obtained using *vg deconstruct* v1.65 for the Minigraph-Cactus, Cactus and PGGB graphs, and using the *minigraph-call* pipeline based on *gfatools bubble* for the Minigraph graph (described in Minigraph documentation with Minigraph v.0.21). The reference genome, used as coordinate system for the output VCF file, was set as GRCh38 for the real human datasets, and the unmutated sequence for the simulated datasets. We ran *vg deconstruct* with the parameter‘-a‘to obtain an exhaustive set of bubbles (without any filter). Notably, we could not use the same bubble caller for all graphs, because vg-deconstruct require P or W lines in GFA format and gfatools-bubbles applies only to rGFA graphs output only by Miningraph. We could not use BubbleGun neither, because it does not report bubbles in VCF format and, more importantly, does not report allelic path traversals within bubbles, which is en essential requirement for annotating the bubbles.

All command lines for graph construction and bubble calling are provided in Supplementary material and in the dedicated public repository https://github.com/ SandraLouise/INVPG annot paper.

### Method for annotating inversion bubbles

We developped INVPG-annot, an automated method to identify, among the bubbles found in a pangenome graph, the ones that correspond to inversion topologies. INVPG-annot takes as input a bubble file in VCF format, as output typically by vg-deconstruct and minigraph-call pipeline, with the following necessary information: the reference and allele sequences, as well as the paths of all alleles of the bubble, given as ordered and oriented lists of nodes (AT field).

The method is composed of three main steps, as shown in Fig. 2. The first step consists in selecting potential inversion bubbles based on the size of their alleles, in order to limit the amount of bubbles to analyze. We select bubbles resembling to balanced SVs by extracting bubbles with at least two alleles such that their length are similar and both greater than 50 bp. We consider two alleles *a*_0_ and *a*_1_, of length *l*_0_ and *l*_1_, to be of similar length if 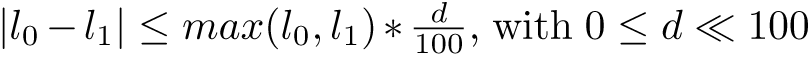 expressed as a percentage. The parameter *d* allows for leniency on the tolerated length difference of inversion alleles, as such difference can originate from innate biological sequence divergence but also from possible computational artefact. In the experiments, we used *d* = 10% for all real and simulated datasets.

In order to annotate inversions among the selected bubbles, we then look for inversion signals in the paths of the bubbles (step 2) and, if that fails, in the alignments of the bubble allele sequences (step 3). For each bubble, all alternative alleles are treated independently and compared to the reference allele.

We consider a path-based inversion signal as a common node between the alternative and reference allele paths that is traversed in opposite ways by the two paths. In order to identify such nodes, we convert the reference and alternative paths into two ordered lists *ord_ref_* = [*x*_1_*,.., x_n_*] and *ord_alt_* = [*y*_1_*,.., y_m_*] of integer node identifiers, which are positive if the node is traversed in forward and negative if the node is traversed in reverse direction. Then for each node identifier *x_i_* in *ord_ref_*, we save *x_i_* as a path-based inversion signal if −*x_i_* is found in *ord_alt_*. For each alternative allele, we calculate a path-based signal score *cov_path_*, which is the fraction of the reference allele’s length covered by path-based inversion signals, with the following formula:

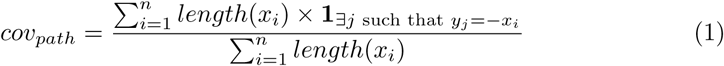

If *cov_path_* is greater than a user-defined threshold (*min_cov_*, with default value of 0.5) for at least one alternative path of the bubble, the bubble is annotated as a path-explicit inversion. If not, we then look for inversion signals in the alignment of the bubble alternative allele sequences to the reference allele. We consider an alignment-based inversion signal as a reverse alignment between the alternative and reference allele sequences. In order to identify alignment-based inversion signals, we align all alternative allele sequences on the reference allele sequence using minimap2 [24] v2.15 with the parameters ‘-cx asm20 –cs-r2k‘. Considering *A* = (*a*_1_*,..a_k_*) the set of reverse alignments between one alternative allele and the reference allele, we calculate the alignment-based signal score *cov_aln_* as:

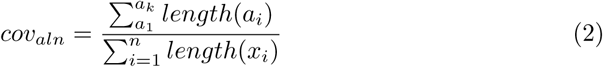

If *cov_aln_*is greater than the *min_cov_* threshold for at least one of the alternative paths, the bubble is annotated as an alignment-rescued inversion.

INVPG-annot is implemented in Python and is available on github (https://github.com/SandraLouise/INVPG_annot). It takes as input a GFA file of the pangenome graph as well as the corresponding VCF containing the bubbles (*vg deconstruct* or *mingraph-call* formats, reporting allele sequences and allele traversals in the graph), and outputs a VCF file containing the set of inversion annotated bubbles along with their topology type in the graph (path-explicit or alignment-rescued). We detail the inversion topology and signal coverage for all alleles of each analyzed bubble using custom tags in the INFO field (INVANNOT and INVCOV).

### Evaluation of inversion detection in the pangenome graphs

To assess the completeness of the sets of annotated inversion bubbles we calculated a recall value as the percentage of known inversions (simulated or real) that were annotated in the graph by at least one of its bubbles. We established the correspondence of the annotated bubbles to the true inversions in the datasets based on their position overlap on the reference genome using bedtools [27] v2.27.1. We considered a given inversion as correctly annotated if a reciprocal overlap of at least 50% was found between the inversion and one of the annotated bubbles (*i.e.* the position of the annotated bubble on the reference covers ≥ 50% of the size of the inversion, and vice-versa), using the options ‘-f 0.5-r‘of the function *bedtools intersect*. We also calculated the number of imprecise inversion bubbles, which we define as annotated bubbles that overlap a true inversion but do not meet the 50% reciprocal overlap requirement. Finally, we calculated the number of redundant inversion bubbles, which we define as an inversion-annotated bubble overlapping at least one other annotated bubbles in the same graph.

## Supplementary information

Supplementary methods and tables are available in Additional file 1.

## Declarations

Ethics approval and consent to participate

Not applicable.

## Consent for publication

Not applicable.

## Availability of data and materials

HPRC, CHM13 and GRCh38 human genome assemblies used in this study can be found at https://data.humanpangenome.org/assemblies. All scripts used to generate simulated datasets and analyze the data are available in the public repository https://github.com/SandraLouise/INVPG annot paper. Simulated datasets support-ing the conclusions of this article are available in the Zenodo repository, at DOI https://doi.org/10.5281/zenodo.18695551. The software INVPG-annot is available on github https://github.com/SandraLouise/INVPG_annot under the open source AGPL license.

## Authors’ contributions

SR, FL and CL conceived and designed the project. SR, SD and CL implemented the scripts and ran the experiments. SR and SD implemented the software. SR, SD, FL and CL wrote the paper. The authors read and approved the final manuscript.

## Acknowledgements

We are thankful to the GenOuest bioinformatics platform for supporting the calcula-tions.

