## Supplementary Material for "Investigating the topological motifs of inversions in pangenome graphs"

### Supplementary material for
### Investigating the topological motifs of inversions in pangenome graphs

Sandra Romain[1,4], Siegfried Dubois[1,2], Fabrice Legeai[1,3], and Claire Lemaitre[1]

[1] *Univ Rennes, Inria, CNRS, IRISA - UMR 6074, F-35000 Rennes, France*
[2] *GenPhySE, Université de Toulouse, INRAE, ENVT, 31326 Castanet Tolosan, France*
[3] *IGEPP, INRAE, Institut Agro, University of Rennes, 35653 Le Rheu, France*
[4] *Current affiliation: Bart's Cancer Institute, Queen Mary University London, EC1M 6AU London, UK*

#### 1 Supplementary material and methods

##### 1.1 Command lines for the different PG pipelines

**Minigraph pipeline**

```
# graph construction
minigraph -cxggs refId.fa simulated.fa > minigraph.gfa

# bubble calling
    #for each haplotype (including the reference)
    minigraph -cxasm --call -t8 minigraph.gfa hapId.fa > hapId-path.bed

ls {*}-path.bed > samples.txt
paste {*}-path.bed | mgutils.js merge -s samples.txt - > merged.bed

mgutils-es6.js merge2vcf -r0 merged.bed > minigraph-bubbles.vcf

# bubble annotation
invpg -v minigraph-bubbles.vcf -g minigraph.gfa -o minigraph_inversions.vcf -m 0.5 -d 10
```

**Minigraph-Cactus pipeline**

```
# Create the pipeline.txt file containing on each line, the haplotype ID
# and the full path of the corresponding sequence fasta file (tab separated)

#Graph construction
JB="jobstore/js.0"
cactus-minigraph $JB pipeline.txt CACTUS/graph.gfa --reference refId \
        --binariesMode singularity
cactus-graphmap $JB pipeline.txt CACTUS/graph.gfa CACTUS/graph.paf  --reference refId \
        --outputFasta CACTUS/graph.sv.gfa.fa.gz --binariesMode singularity
cactus-align $JB pipeline.txt CACTUS/graph.paf CACTUS/graph.hal --pangenome --outGFA \
        --outVG --reference refId --workDir CACTUS/workdir" --binariesMode singularity
cactus-graphmap-join $JB --vg CACTUS/graph.vg --outDir CACTUS/outdir" --outName "final" \
        --reference refId --clip 0 --filter 0 --binariesMode singularity

gzip -d CACTUS/outdir/final.full.gfa.gz

# Convert the graph in GFA1.1 to GFA1.0 format with all P-lines
```

```
vg convert -g -f -W CACTUS/outdir/final.full.gfa > mgc.gfa

# bubble calling
vg deconstruct -a mgc.gfa -p refId > mgc-bubbles.vcf

# bubble annotation
invpg -v mgc-bubbles.vcf -g mgc.gfa -o mgc_inversions.vcf -m 0.5 -d 10
```

##### PGGB pipeline

```
# prepare input haplotypes
cat haplotype1.fa haplotype2.fa > all_haplotypes.fa

# graph construction
pggb -i all_haplotypes.fa -o outdir -n 2 -t 8 -p 90 -s 5k -V refId
mv outdir/all_haplotypes.fa.*.smooth.final.gfa pggb.gfa

# bubble calling
vg deconstruct -a pggb.gfa -p refId > pggb-bubbles.vcf

# bubble annotation
invpg -v pggb-bubbles.vcf -g pggb.gfa -o pggb_inversions.vcf -m 0.5 -d 10
```

##### Cactus pipeline

```
# Create the input_cactus.txt file
# First line of this file = binarized genome tree
# Then one line per haplotype, with its ID and the absolute path of the corresponding
# sequence fasta file (tab separated)
    Eg.
    (refId:1.0,simulated:1.0);
    refId   /path/to/refId.fa
    simulated   /path/to/simulated.fa

#Graph construction
cactus --binariesMode singularity ./js input_cactus.txt cactus.hal
hal2vg --noAncestors --refGenomes refId cactus.hal > cactus.pg
vg convert --gfa-out -W cactus.pg > cactus.gfa

# bubble calling
vg deconstruct -a cactus.gfa -p refId > cactus-bubbles.vcf

# bubble annotation
invpg -v cactus-bubbles.vcf -g cactus.gfa -o cactus_inversions.vcf -m 0.5 -d 10
```

### 2 Supplementary tables and figures

#### 2.1 Details on inversion annotations for the different PG pipelines on the 2-haplotypes simulated datasets

Table 1: Inversion bubble statistics of the pangenome graphs obtained for the 2-haplotype simulated genome sets with 100 inversions and with varying percentages of punctual polymorphism, for each pangenome graph constructing tool used. The number of inversion bubbles indicates the total number of bubbles annotated as inversions by our method. The "Non redundant and TP bubbles" column indicates the number of inversion bubbles that do not overlap any other inversion bubble and that are True Positive bubbles (ie. with at least 50 % reciprocal overlap with a simulated inversion). The "Redundant bubbles" column indicates the number of inversion bubbles that overlap at least one other inversion bubble and all overlap a simulated inversion by at least one bp. "Imprecise" bubbles indicate the number of inversion bubble that overlap a simulated inversion by less than 50 % (not counted in the Recall). The Recall column indicates the percentage of simulated inversions (among the 100) that have a significant reciprocal overlap with at least one inversion annotated bubble. ("MC": Minigraph-Cactus).

| Genome set | PG pipeline | # Inversion bubbles | # Non redundant TP bubbles | # Redundant bubbles | # Imprecise bubbles | Recall (%) |
|---|---|---|---|---|---|---|
| 0% SNP | Cactus | 92 | 92 | 0 | 0 | 92 |
| | Minigraph | 7 | 4 | 0 | 3 | 4 |
| | MC | 28 | 16 | 2 | 5 | 16 |
| | PGGB | 92 | 90 | 0 | 2 | 90 |
| 0.1% SNP | Cactus | 92 | 92 | 0 | 0 | 92 |
| | Minigraph | 75 | 72 | 0 | 3 | 72 |
| | MC | 90 | 78 | 0 | 4 | 78 |
| | PGGB | 90 | 89 | 0 | 1 | 89 |
| 1% SNP | Cactus | 92 | 92 | 0 | 0 | 92 |
| | Minigraph | 75 | 75 | 0 | 0 | 75 |
| | MC | 92 | 82 | 0 | 2 | 82 |
| | PGGB | 90 | 89 | 0 | 1 | 89 |
| 5% SNP | Cactus | 91 | 67 | 24 | 1 | 90 |
| | Minigraph | 73 | 73 | 0 | 0 | 73 |
| | MC | 85 | 79 | 0 | 3 | 79 |
| | PGGB | 89 | 89 | 0 | 0 | 89 |

#### 2.2 Details on inversion annotations for the different PG pipelines on the 10-haplotypes simulated datasets

Table 2: Inversion bubble statistics of the pangenome graphs obtained for the 10-haplotype simulated genome sets with 100 inversions and with 1% of punctual polymorphism, for each pangenome graph constructing tool used. The number of inversion bubbles indicates the total number of bubbles annotated as inversions by our method. The "Non redundant TP bubbles" column indicates the number of inversion bubbles that do not overlap any other inversion bubble and that are True Positive bubbles (ie. with at least 50 % reciprocal overlap with a simulated inversion). The "Redundant bubbles" column indicates the number of inversion bubbles that overlap at least one other inversion bubble and all overlap a simulated inversion by at least one bp. "Imprecise" bubbles indicate the number of inversion bubble that overlap a simulated inversion by less than 50 % (not counted in the Recall). The Recall column indicates the percentage of simulated inversions (among the 100) that have a significant reciprocal overlap with at least one inversion annotated bubble. ("MC": Minigraph-Cactus).

| Genome set | PG pipeline | # Inversion bubbles | # Non redundant TP bubbles | # Redundant bubbles | # Imprecise bubbles | Recall (%) |
|---|---|---|---|---|---|---|
| 1% SNP | Cactus | 162 | 30 | 132 | 1 | 90 |
| | Minigraph | 76 | 76 | 0 | 0 | 76 |
| | MC | 88 | 85 | 2 | 8 | 86 |
| | PGGB | 94 | 92 | 0 | 2 | 92 |

#### 2.3 Performances of INVPG-annot on the real human pangenome graphs

Table 3: Cumulated graph size, total number of bubbles and running time of INVPG-annot on the human chromsome 7 and chromosome X pangenome graphs produced by the 4 pipelines. INVPG-annot was run on a single core of Intel(R) Xeon(R) CPU E5-2630 v4 @ 2.20GHz. "MC": Minigraph-Cactus.

| PG pipeline | Graph size (Mbp) | # Bubbles | Running time (min) | RAM (Mb) |
|---|---|---|---|---|
| Cactus | 343 | 1,512,364 | 8min07s | 669 |
| Minigraph | 319 | 4,620 | 2min12s | 174 |
| MC | 371 | 1,042,944 | 14min08s | 442 |
| PGGB | 310 | 1,024,316 | 35min11s | 535 |
